# Neural sensitivity to the heartbeat is modulated by fluctuations in affective arousal during spontaneous thought

**DOI:** 10.1101/2025.03.26.645574

**Authors:** David Braun, Lotus Shareef-Trudeau, Swetha Rao, Christine Chesebrough, Julia W. Y. Kam, Aaron Kucyi

**Affiliations:** Department of Psychological & Brain Sciences, Drexel University, 19104; Northwell Health Feinstein Institutes for Medical Research, 11030; Department of Psychology, University of Calgary, T2N 1N4

## Abstract

Spontaneous thoughts, occupying much of one’s awake time in daily life, are often colored by emotional qualities. While spontaneous thoughts have been associated with various neural correlates, the relationship between subjective qualities of ongoing experiences and the brain’s sensitivity to bodily signals (i.e., interoception) remains largely unexplored. Given the well-established role of interoception in emotion, clarifying this relationship may elucidate how processes relevant to mental health, such as arousal and anxiety, are regulated. We used EEG and ECG to measure the heartbeat evoked potential (HEP), an index of interoceptive processing, while 51 adult participants (34 male, 20 female) visually fixated on a cross image and let their minds wander freely. At pseudo-random intervals, participants reported their momentary level of arousal. This measure of affective arousal was highly variable within and between individuals but was statistically unrelated to several markers of physiological arousal, including heart rate, heart rate variability, time on task, and EEG alpha power at posterior electrodes. A cluster-based permutation analysis revealed that the HEP amplitude was increased during low relative to high affective arousal in a set of frontal electrodes during the 340 – 356 millisecond window after heartbeat onset. This HEP effect was more pronounced in individuals who reported high, relative to low, levels of trait anxiety. Together, our results offer novel evidence that at varying levels of trait anxiety, the brain differentially modulates sensitivity to bodily signals in coordination with the momentary, spontaneous experience of affective arousal—a mechanism that may operate independently of physiological arousal.

**Significance Statement:** Our findings highlight the relationships between spontaneous fluctuations in affective arousal, brain-body interactions, and anxiety, offering new insights into how interoception fluctuates with changes in internal states. By showing that interoceptive processing is heightened during lower affective arousal and that this effect is amplified in individuals with higher trait anxiety, our study suggests the brain adaptively downregulates interoceptive sensitivity in response to fluctuating internal states. These results have implications for understanding how spontaneous thoughts shape interoception and emotion, particularly in clinical contexts where dysregulated interoception is linked to anxiety and mood disorders. More broadly, our work underscores the need to distinguish between different forms of arousal, advancing understanding of the taxonomy and ways of measuring arousal.

## Introduction

The human brain continually processes signals from the body to guide perception and behavior, a capacity referred to as *interoception* (Engelen et al., 2023). Although early theorists (e.g., James, 1884/1948; Sherrington, 1906) noted its role in cognition and emotion, interoception has recently become a critical focus in cognitive neuroscience (Azzalini et al., 2019; Kluger et al., 2024). A method of measuring interoception—first introduced in the late 20^th^ century(Montoya et al., 1993; Schandry et al., 1986) and now commonly applied with refined approaches (Park & Blanke, 2019)—is the *heartbeat-evoked potential (HEP)*, which involves averaging neural electrophysiological signals time-locked to features of the electrocardiogram (*ECG*) representing the neural processing of heartbeats. Larger HEP amplitude is often interpreted as greater interoception (Judah et al., 2018; Pang et al., 2019; Verdonk et al., 2024), and the HEP has been linked to affective arousal (Fourcade et al., 2024), a factor central to the present investigation.

Arousal is historically defined as physiological and psychological activation or intensity (Russell, 1980; Russell & Barrett, 1999). Recently, researchers have debated whether arousal is best characterized as unidimensional or multidimensional (Sabat et al., 2025), for example separating affective and physiological aspects as independent subcomponents (Reid et al., 2024; Satpute et al., 2019). Differences in the HEP have been found in frontal electrodes in the 200–450 ms interval post-heartbeat across various conditions that broadly targeted arousal, including exposure to emotional stimuli (Marshall et al., 2017, 2018, 2020; Sel et al., 2017), food deprivation (Schulz et al., 2015), pain (Shao et al., 2011), and cortisol injections (Schulz et al., 2013). Notably, stimulus- and task-evoked arousal has previously been associated with both increased (Luft & Bhattacharya, 2015) and diminished (Fourcade et al., 2024) interoception.

Another way to target arousal is to measure natural, spontaneous fluctuations during wakeful rest (Chang et al., 2016; Kucyi & Parvizi, 2020), an approach that is also increasingly used to investigate the neural basis of ongoing mental experience (e.g., “spontaneous thoughts” or “mind-wandering”; Christoff et al., 2016; Kam et al., 2021; Shareef-Trudeau et al., In press; Smallwood et al., 2018). For example, HEP amplitudes vary depending on spontaneous fluctuations in self- or other-referenced thoughts (Babo-Rebelo et al., 2019; Babo-Rebelo, Richter, et al., 2016; Babo-Rebelo, Wolpert, et al., 2016). However, no study to our knowledge has linked the HEP to spontaneous fluctuations in affective arousal. Spontaneous mental experiences are pervasive in daily life (Killingsworth & Gilbert, 2010), and the arousal component of experience is particularly relevant to mental health conditions involving anxiety.

Anxiety can involve dysregulated interoception, such as heightened or inconsistent sensitivity to bodily signals (Khalsa & Lapidus, 2016). Individuals scoring higher in anxiety tend to exhibit larger HEP amplitudes (Judah et al., 2018; Pang et al., 2019; Verdonk et al., 2024). Given the intimate associations between (i) anxiety and interoception (Janelle, 2002; Noteboom et al., 2001; Roos et al., 2021) and (ii) anxiety and abnormal spontaneous thinking (Fell et al., 2023), spontaneous fluctuations in affective arousal and their interoceptive underpinnings may prove important for understanding anxiety.

Here we examined the relationship between subjective qualities of ongoing experiences and interoception as well as how this relationship might be influenced by anxiety. Participants completed anxiety surveys and underwent EEG and ECG recordings during a wakeful resting state, intermittently rating affective arousal via experience sampling. To better understand our “affective arousal” construct, we assessed whether self-reported arousal converged with physiological measures or was linked to other aspects of ongoing experience. We next leveraged HEP analysis to examine whether affective arousal is associated with higher or lower levels of interoceptive processing. Finally, we explored whether individual variability in trait anxiety modulated the relationship between HEP amplitude and affective arousal. Our findings offer important implications for how arousal is conceptualized and how spontaneous internal fluctuations shape the link between interoception and emotion.

## Materials and Methods

### Participants

Fifty-four participants (34 female, 20 male) were recruited to participate in this study. Participants were between the ages of 18 and 31, and 50.00% identified as Asian, 35.19% identified as White, 11.11% identified as Black or African American, 1.85% identified as two or more races, and 1.85% identified as other. Participants were recruited both from undergraduate psychology courses for optional course credit and from recruitment posters placed around Drexel and Temple University campuses. Those recruited through the posters received $60 USD of compensation for participation. All participants were fluent in English, had normal or corrected-to-normal vision, were right-handed, and reported no history or current diagnosis of any neurological or psychiatric disorders. All participants provided informed consent for procedures approved by the Drexel University Institutional Review Board. One participant was excluded due to removal of an excessive number of noisy epochs in EEG recordings, one participant was excluded due to EEG recording failure, and one participant was excluded due to behavioral data recording failure, leaving 51 participants for the final analysis.

### EEG Data Acquisition

EEG recordings were obtained using a 32-channel Brain Products system, including the BrainAmp MRPlus and the BrainAmp ExG (*BrainCap MR*, 2020). We recorded EEG data with the BrainVision Recorder Vers. 1.25.0201 on a Windows operating system using a Dell Latitude 7430 64-bit operating system with a 12^th^ Gen Intel Core i7-1265U and 16GB of RAM. Data were continuously sampled at 5000 Hz. We fitted participants using the BrainCap MR (based on the extended 10/20 system) with embedded carbon wire loops. The system includes ground and reference electrodes, each with 15 kOhms of resistance, 31 scalp electrodes with 10 kOhms of resistance each and an ECG channel with 20 kOhms of resistance. We recorded heartbeat activity via the ECG which was applied to the center of each participant’s back as far down as the cord could reach, typically mid-back (Yamashita et al., 2024). Cap setup was concluded when all electrodes’ impedance measurements were at levels below 20 kOhms.

### Mental Health Surveys

Prior to EEG setup, participants filled out a series of surveys pertaining to their demographic (e.g., age, sex, race) and mental health status. Of these, only the Generalized Anxiety Disorder-7 (GAD-7; Spitzer et al., 2006) and the state scales of the State Trait Anxiety Inventory (STAI-S; Speilberger et al., 1983) were analyzed to capture both trait and state anxiety, respectively, given our focus on the links of anxiety with arousal and interoception (for the complete set of surveys administered in the experiment, see *Table S1*). The state anxiety measure (STAI-S) was a late addition to the experiment after 23 participants’ data had been collected. All surveys, except for the STAI-S, were administered online using REDCap electronic data capture tools (Harris et al., 2009, 2019). STAI-S was collected either online or with paper and pencil. Participants were asked to complete all but the state anxiety (i.e., STAI-S) measure prior to arriving for the experiment. Both of the GAD-7 and STAI-S scales have shown high validity and reliability across a wide range of contexts and populations (Gunning et al., 2010; Löwe et al., 2008; Pretorius & Padmanabhanunni, 2023; Vitasari et al., 2011). The GAD-7 and STAI-S are shown to have high internal consistency, both in past research (Cronbach’s α*_GAD_* = 0.89, Löwe et al., 2008; α_STAI_ = 0.87, Spielberger et al., 2017) and in our sample (α*_GAD_* = 0.91, α*_STAI_* = 0.89).

### Experience Sampling

Throughout the EEG/ECG experiment, participants were intermittently prompted to report on the nature of their ongoing experience using thought probes. Each probe consisted of 13 experience sampling items presented sequentially in a fixed order. For each item, participants provided a rating on a 0–100 slider scale assessing a different dimension of their experience in the moments immediately preceding the probe’s onset. The primary experience sampling item relevant to this study measured affective arousal, asking: *“How activated or energized were you feeling?”* The highest end of the anchor was labeled *“Completely activated”*, while the lowest end was labeled *“Completely deactivated.”* Affective arousal was defined to participants as *“feelings of energy linked to emotion, which may result in heightened bodily energy, tension, or responsiveness”,* a definition which is consistent with prior research (Barrett, 1998; Bliss-Moreau et al., 2020; Haj-Ali et al., 2020; Kron et al., 2015; Kuppens et al., 2013; Maher et al., 2019; Nook et al., 2017; Russell, 1980, 2003; Russell & Barrett, 1999). Other relevant experience sampling items for present purposes included probes about future thinking (“Were your thoughts oriented towards the future?” from “Not future oriented” to “Completely future oriented”), deliberate thinking (“How intentional were your thoughts?” from “Completely unintentional” to “Completely intentional”), self-related thinking (“Were your thoughts about yourself?” from “Nothing about you” to “Completely about you “), and difficulty to disengage from thinking (“How difficult was it to disengage from your thoughts?” from “Extremely easy” to “Extremely difficult”). Participants additionally rated their level of confidence in their responses at the end of each thought probe (from “Completely confident” to “Completely unconfident”). For the complete set of experience sampling items used in the experiment, see *Table S2*.

## Procedure

Upon arrival, participants provided informed consent and were fitted with an EEG cap while receiving task instructions. They were introduced to each experience sampling item and tested for comprehension using a hypothetical scenario in which a student was experiencing stress while preparing for an exam, and participants were asked to respond to experience sampling items as if they were the hypothetical student studying for the exam. The experimenter provided feedback to ensure participant comprehension of instructions before proceeding, and evidence of comprehension was supported by participants giving generally high ratings of confidence in their responses throughout the experiment (*M =* 17.89, where 0 is most confident and 100 is least; *SD* = 14.41).

The main task consisted of experience sampling during wakeful rest (see *Figure 1A*). During wakeful rest, participants sat at a computer and were instructed to “Let your mind wander freely” while focusing on a black screen with a central white fixation cross. At a randomized integer interval every 15–45 s, the fixation cross was replaced by a thought probe with a series of experience sampling items, which participants answered using arrow keys on a laptop computer to adjust a slider scale and entered their chosen response using the spacebar. After the last item, the fixation cross reappeared, and the cycle repeated.

**Figure 1.**
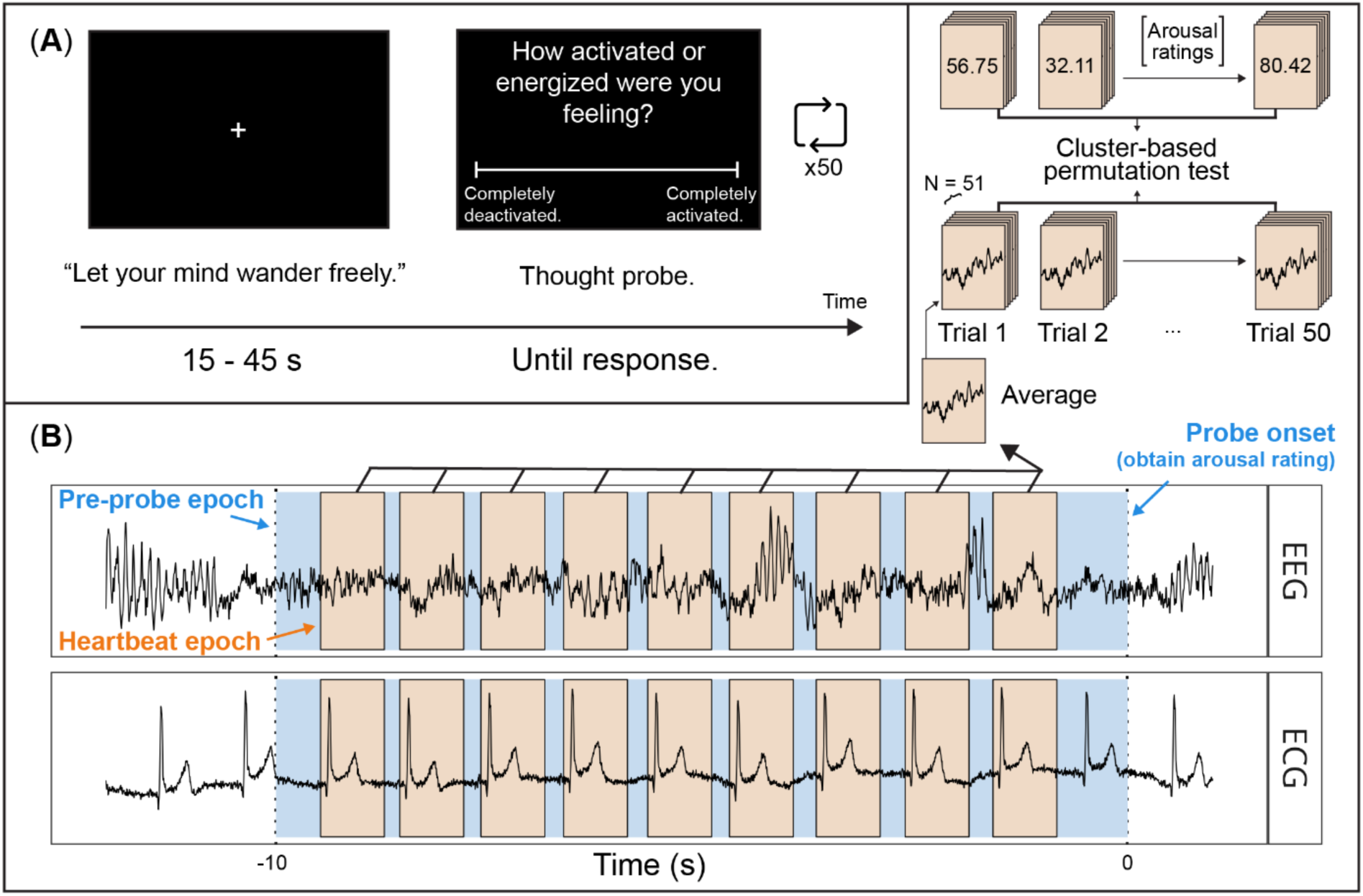
Trial sequence and EEG/ECG analysis approach. (A) Participants were instructed to “Let your mind wander freely” while focusing on a black screen with a central, white fixation cross. EEG and ECG were simultaneously recorded throughout. After 15 – 45 s, a series of thought probes appeared, prompting participants for ratings about their experience. This trial sequence was repeated 50 times throughout the experiment. (B) Pre-probe epochs were first defined in the window 10 s prior to thought probe onset. Within pre-probe epochs, heartbeat epochs were defined in the EEG data from −0.1 – 0.65 s around each R-peak in the ECG data. We then conducted trial-wise regressions, using EEG voltage at each timepoint and channel within the heartbeat-locked epochs to predict affective arousal ratings. This yielded a timepoint-by-channel map of regression slopes for each participant, which we submitted to a cluster-based permutation analysis using one-sample *t*-tests against zero to identify significant clusters.

Prior to the wakeful rest and experience sampling task, participants first alternated between opening and closing their eyes for eight minutes to validate that EEG was recording as expected. Participants next performed the main phase of the experiment, where each cycle of wakeful rest and experience sampling repeated 10 times per run, with 5 total runs, yielding 50 total trials (i.e., thought probe sets). Prior to the start of each task run, the experimenter exited the room. The experimenter reentered the room between each run to give the participants a short break and cue the next task/run. Data collection lasted approximately 60 minutes, with the full session taking about 2 hours.

The first 22 participants were tested in a windowless lab room, which was equipped with a stimulus laptop for task administration and an EEG recording laptop. During the main task, the experimenter visually monitored participants remotely via Zoom (*Zoom Video Communications*, 2024) using a separate monitoring computer. Due to equipment and facility constraints, the remaining 31 participants were tested in a different room that included a glass wall. Participants were seated with their backs to the glass wall and, to minimize external distractions during the task, participants wore earplugs.

## EEG Preprocessing

Electrophysiological data preprocessing was performed using custom Python scripts based on the MNE library (Gramfort et al., 2013; Larson et al., 2023). Data were downsampled to 250 Hz, bandpass filtered between 1 and 55 Hz, and referenced to the common average. Initial artifact removal was conducted using Extended Infomax independent components analysis (ICA; Lee et al., 1999), with automatic classification and rejection of non-brain artifacts via ICLabel (Pion-Tonachini et al., 2019). Noisy channels were identified via manual inspection and interpolated on the pre-ICA data. Five participants had one channel interpolated, two participants had two channels interpolated, and two participants had three or four channels interpolated. A second ICA decomposition was then performed, and ICLabel results were manually reviewed for further artifact rejection. We chose not to perform baseline correction for HEP analyses because, unlike in stimulus-evoked designs, heartbeat-evoked designs lack a true baseline. Cardiac activity is cyclical, and electrical activity before a heartbeat can reflect both the preceding heartbeat and preparatory processes for the upcoming one (Azzalini et al., 2019).

For HEP analyses, R-peaks in the ECG signal were detected using MNE’s preprocessing library. We defined the 10 s period prior to the onset of the thought probe as the time period of interest, as brain states linked to self-report probes can emerge up to 10 s before probe onset (Christoff et al., 2009; Kucyi et al., 2024). Heartbeats occurring within 10 s prior to each thought probe onset were retained for analysis. Following prior work (Petzschner et al., 2019), heartbeats within 700 ms of each other were excluded to prevent overlapping epochs. Within this 10 s pre-probe period, epochs were defined on the EEG channels from −100 ms to +650 ms relative to each R-peak. Noisy epochs were automatically identified with the autoreject library (Jas et al., 2017), and all epochs were manually inspected before removal.

The cardiac field artifact (CFA) is noise introduced into the EEG signal due to electrical activity from the heart. While ICA is a powerful method for removing many types of artifacts, including the CFA, from EEG (Debener et al., 2010; Devuyst et al., 2008; Hoffmann & Falkenstein, 2008), no known method can entirely remove the CFA from EEG data. To ensure the CFA was not impacting the EEG signal differently across experimental conditions, we conducted several control analyses. First, we examined the association between ECG metrics—including peak and mean ECG amplitude—and affective arousal (Petzschner et al., 2019). This association was examined by first averaging both ECG metrics in the significant time window revealed by the cluster-based permutation on the EEG data (see below), and then separately regressing both averaged ECG metrics on to affective arousal ratings across trials, using Bayesian estimation to provide Bayes factors in support of the null hypothesis. Second, we applied the same cluster-based permutation analysis used in our EEG analysis (see below) directly to the ECG data. Any statistically significant associations between ECG amplitude and affective arousal would raise concerns about a potential CFA-related confound.

For spectral analyses involving parieto-occipital alpha power, epochs were defined over the full 10 s period prior to thought probe onset. Noisy epochs were again removed by manually inspecting results from the autoreject library. We examined parieto-occipital alpha power in the 10 s pre-probe period, as past studies have shown a connection between parieto-occipital alpha power and various types of arousal (Hofmann et al., 2021; Lozano-Soldevilla, 2018; Schubring & Schupp, 2021; Tuladhar et al., 2007). To examine parieto-occipital power, the EEG signal was decomposed into time-frequency dimensions via MNE’s Morlet wavelet transform algorithm with 50 logarithmically spaced frequencies from 2 to 30 Hz and a variable number of cycles per wavelet (*n*_cycles_ = freq/2) for occipital and parietal channels only, namely O1, O2, Oz, P3, P4, P7, P8, and Pz (Fourcade et al., 2024). We decomposed the signal from 2-30 Hz to allow for exploratory flexibility in examining power across multiple frequency bands, though our primary focus in this study is on parieto-occipital alpha power.

## Experimental Design and Statistical Analysis

### Affective Arousal, Physiological Arousal, and Other Experience Sampling Measures

To assess the relationship between physiological and affective arousal, measures of the two constructs were compared. Measures of physiological arousal included heart rate, heart rate variability (HRV), time-on-task, and parieto-occipital alpha power amplitude (Hofmann et al., 2021; Lozano-Soldevilla, 2018; Schubring & Schupp, 2021; Tuladhar et al., 2007), each of which except for time-on-task were summarized over all heartbeats in the 10 s period prior to probe onset (i.e., not removing heartbeats occurring within 700 ms of each other, a procedure only done for the HEP analyses). Time-on-task, calculated as the probe (i.e., trial) order number, was included because of its well established links to self-reported fatigue, impaired task performance, and negative association with physiological arousal measures (Gergelyfi et al., 2015; Herlambang et al., 2019; Karthikeyan et al., 2021, 2022; Matuz et al., 2019; Melo et al., 2017; Qin et al., 2021; Shi et al., 2022; Zhao et al., 2012). HRV was calculated using the root mean square of differences between successive heartbeats (Shaffer & Ginsberg, 2017). Affective arousal was measured by self-reported levels of arousal obtained at each thought probe.

We used hierarchical Bayesian linear regressions to estimate relationships between physiological and affective measures of arousal. These regressions were implemented via the brms (Bürkner, 2017) and Stan (Stan Development Team, 2024) packages in R (R Core Team, 2024) using default, weakly-informative priors. Hierarchical Bayesian estimation is ideal for accurately estimating trends at both the group and individual level. Separate models were fit for each physiological arousal measure—heart rate, HRV, time on task, and alpha power—with the physiological arousal measure predicting affective arousal. Random intercepts and slopes were included into the model, grouped by participant. To aid estimation, affective arousal was z-scored within participants and alpha power was log-transformed prior to model sampling. Estimation of each model used four chains, each drawing 5,000 samples (including 2,500 warm-up). Convergence was assessed by inspecting *R̂* values, ensuring all values were ≤ 1.01. All models showed adequate effective sample sizes and no convergence warnings. We summarized both group- and subject-level slope estimates using posterior means and 95% credible intervals. Intervals excluding zero were interpreted as evidence of a nonzero relationship. These posterior summaries were complemented by significance estimates from traditional (i.e., frequentist) hierarchical models estimated using *lme4* (Bates et al., 2015) using the Satterthwaite degrees of freedom approximation from the *lmerTest* package (Kuznetsova et al., 2017) to obtain *p* values. We used the same modeling methods to assess the degree to which ratings of affective arousal were related to other self-reported experience sampling measures.

### Cardiac Interoceptive Sensitivity and Affective Arousal

Cardiac interoceptive sensitivity was analyzed in relation to affective arousal by first averaging heartbeat epochs at the trial level within participants. Time points in the epochs less than 250 ms post-R peak were excluded from statistical analysis to avoid overlap with the CFA (Al et al., 2021; Fourcade et al., 2024; Tanaka et al., 2023). Moreover, much previous research using both scalp and intracranial EEG have found HEP effects prior to 450 ms post heartbeat (Al et al., 2020, 2021; Babo-Rebelo, Wolpert, et al., 2016; Canales-Johnson et al., 2015; Flasbeck et al., 2024; Fukushima et al., 2011; Kern et al., 2013; Park et al., 2018; Yuan et al., 2008), including those studies focused specifically on arousal (Coll et al., 2021; Fourcade et al., 2024). Thus, in line with prior work, we chose to restrict our statistical analysis to the 250 – 450 ms window post heartbeat. For each participant, we performed a regression at every channel and epoch timepoint, using affective arousal ratings to predict EEG voltage across trials. The resulting slope coefficients were compiled across participants, yielding a time (250–450 ms) by channel (31 EEG electrodes) array of slopes per participant. We then submitted these slope arrays to a group-level cluster-based permutation test, with the statistical test being a one-sample t-test against zero, identifying clusters where EEG voltage showed a consistent linear relationship with arousal ratings across participants. The cluster-based permutation test used a *p* = .01 cluster forming threshold, 5,000 simulations, and evaluated two-tailed significance.

#### Validating Heartbeat Evoked Effects

To validate that effects observed in cluster-based permutation analyses were indeed heartbeat-evoked—and not driven by external events or spontaneous drift unrelated to heartbeats—we conducted a *surrogate heartbeat control analysis* (Azzalini et al., 2019; Babo-Rebelo, Wolpert, et al., 2016; Park et al., 2016; Park & Blanke, 2019; Steinfath et al., 2024). This surrogate analysis compares the observed cluster effect to a null distribution generated by disrupting the time-locking between heartbeats and EEG epochs. The test statistic for the surrogate analysis was derived by performing a participant-specific median-split version of the cluster-based permutation test described above (as the nature of repeated simulations required for the surrogate analysis rendered the regression method computationally infeasible). Trial-level epochs were further averaged to a within-participants median split on affective arousal ratings, generating a time (250–450 ms) by channel (31 EEG electrodes) array of EEG voltages for each affective arousal condition (i.e., high and low) for each participant. The statistical test was a paired *t* test, comparing these arrays across conditions and participants. Obtaining the test statistic for the surrogate test involved summing these *t* values from the significant cluster. To generate the null distribution, we simulated surrogate heartbeats by randomly sampling *k* timepoints—where *k* is equal to the number of true heartbeats—within each 10 s pre-probe period, continuing to sample if any surrogate heartbeat times were in the set of true heartbeat times. We constructed EEG epochs (−100 to +650 ms relative to each surrogate timepoint) and applied the autoreject library (Jas et al., 2017) to identify and remove noisy epochs. Autoreject was fit separately for each participant across all epochs, using its automated thresholding and sensor interpolation methods to optimize artifact rejection without manual intervention. After cleaning, the remaining epochs were processed using the same median-split pipeline as in the main analysis, resulting in a time (250–450 ms) by channel (31 electrodes) array for each condition and participant. We then submitted these arrays to a cluster-based permutation analysis identical to the original, summed the *t* values within each cluster, identified the cluster with the largest absolute value *t* sum, and recorded the original (signed) sum for that cluster as one observation in the null distribution. This process was repeated 100 times, yielding an empirical null distribution of cluster statistics. The original test statistic was evaluated for two-tailed significance by comparing its percentile within this null distribution (i.e., whether it exceeded the 97.5^th^ or fell below the 2.5^th^ percentile).

### Cardiac Interoceptive Sensitivity, Affective Arousal, and Anxiety

To examine whether individual differences in anxiety modulates the link between cardiac interoceptive sensitivity and affective arousal, the magnitude of HEP effects was correlated with anxiety scale scores. To determine HEP effect magnitude for each participant, we averaged the regression slopes in a significant time by channels cluster. This averaged slope was correlated separately with GAD (trait anxiety) and STAI-S (state anxiety) composite scores.

For *t* tests and non-hierarchical regressions reported in the results, we report Bayes factors characterizing evidence either in favor of or against the null hypothesis using the *BayesFactor* package (Morey et al., 2011; Morey & Rouder, 2011; Rouder et al., 2009) in the *R* programming language (R Core Team, 2024). We use the conventional notation of *BF_01_* to denote evidence in favor of the null hypothesis and *BF_10_* to denote evidence in favor of the alternative hypothesis. We used default prior settings in all applications, which are a noninformative Jeffrey’s prior placed on the variance of the normal population, and a Cauchy prior placed on the standardized effect size. We interpret Bayes factors using canonical thresholds: < 3 as anecdotal evidence, > 3 & < 10 as substantial evidence, > 10 & < 31.6 as strong evidence, > 31.6 & < 100 as very strong evidence, and > 100 as decisive evidence (Jeffreys, 1961/1998). Note that for hierarchical Bayesian models, we do not report Bayes factors, as estimating marginal likelihoods necessary for these Bayes factors is notoriously difficult. Estimation methods such as bridge sampling, while available, are known to be highly sensitive to prior specification and model complexity (Gronau et al., 2017, 2020).

## Results

### Sufficient trial-level data and no modulation of cardiac field artifact by affective arousal

Given that the HEP is relative to the onset of a heartbeat and not to an external stimulus, enough time needs to elapse between heartbeats to cleanly measure the HEP (Petzschner et al., 2019). To prevent overlapping heartbeat events, we dropped detected R peaks that were within 700 ms of each other, resulting in an average dropout rate of 11.48% (*SD* = 14.28%) per participant. An average of 9.76 epochs per trial (i.e., 10-second pre-thought probe interval) (*SE* = 0.20) were retained per participant, and there was strong evidence that the number of epochs retained per trial was not associated with affective arousal ratings (*b* = 0.001, *CIs* = [−0.002,0.002], *t*(2355) = 0.43, *p* = .666, *BF*_01_ = 19.68). These results suggest that a sufficient number of epochs were being averaged per trial.

To control for the CFA, we sought to ensure that peak and mean ECG amplitude were not associated with affective arousal ratings. Comparing peak ECG amplitude within the significant time window (revealed by the cluster-based permutation analysis, 340 – 356 ms post R-peak; see below) also revealed strong evidence in favor of no association between peak ECG amplitude and affective arousal ratings (*b* = −1.12 × 10^-8^, 95 *CIs* = [−1.08 × 10^-7^, 8.50 × 10^-8^], *t*(2355) = 0.236, *p* = .819, *BF*_01_ = 21.03). Additionally, there was strong evidence in favor of no association between difference in mean ECG amplitude and affective arousal ratings (*b* = −5.43 × 10^-9^, 95 *CIs* = [−9.82 × 10^-8^, 8.74 × 10^-8^], *t*(2355) = 0.115, *p* = .909, *BF*_01_ = 21.44). Further, a cluster-based permutation analysis of ECG signals, using the same parameters as the EEG analysis (see below), failed to find any significant clusters. These results suggest that the CFA was well controlled and not modulated by affective arousal. Finally, a visual inspection revealed that the averaged ECG waveform was nearly identical across a participant-specific median split in affective arousal ratings (*Figure S1*).

### Affective arousal is distinct from physiological arousal and other experience sampling measures

We next examined how affective arousal fluctuates within and between participants and how it relates to other measures collected in the experiment. Affective arousal, rated on a 100-point scale, varied substantially between participants (*M* = 49.1, *SD* = 12.4), with some participants exhibiting a bimodal pattern of responding while others exhibited a unimodal, normal-shaped pattern (see *Figure 2A and Figure S2*). For some participants, affective arousal varied widely over time, whereas others displayed more stable patterns (see *Figure 2B*).

**Figure 2.**
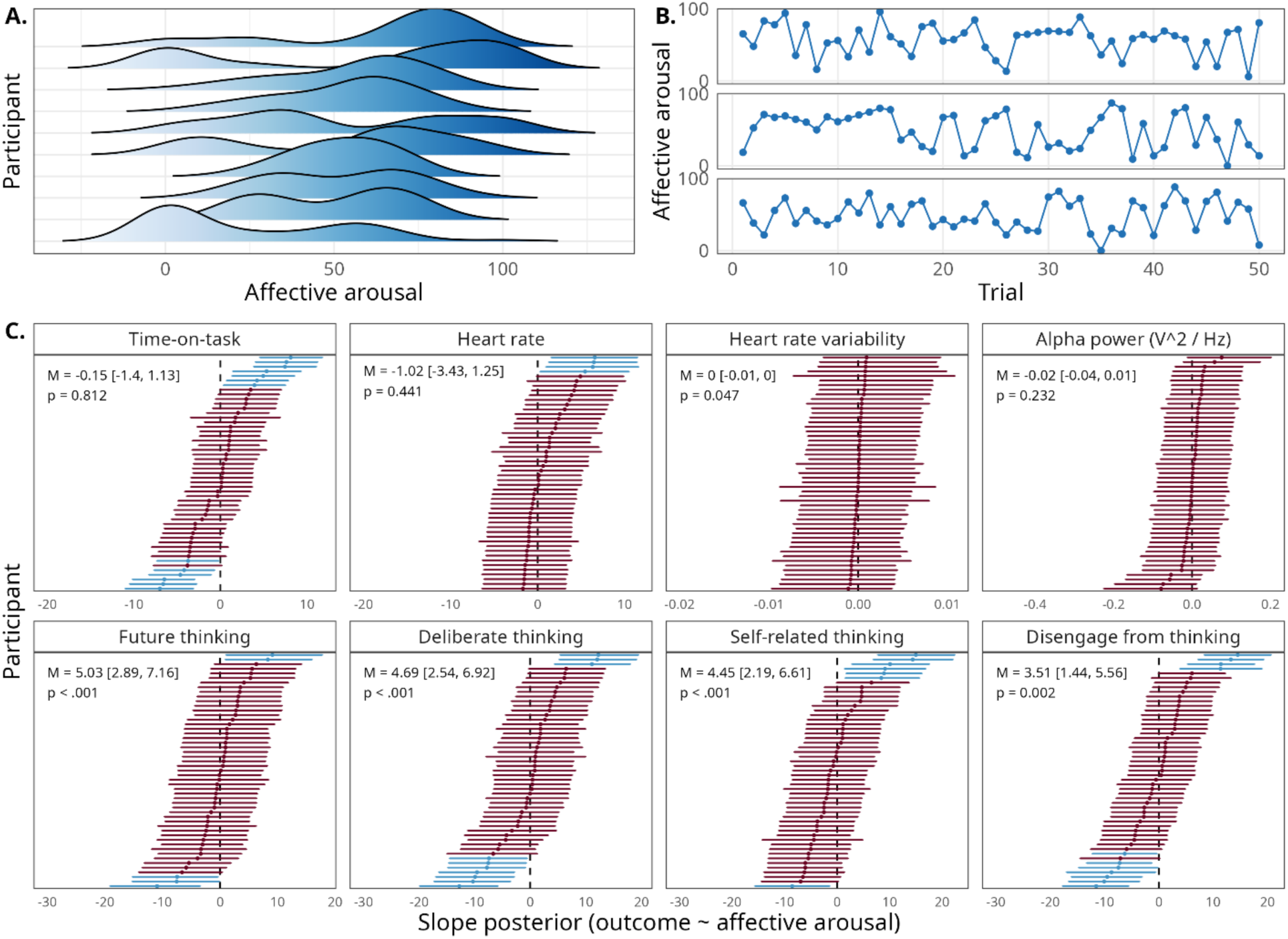
Characteristics of affective arousal ratings and relationships with other physiological and subjective measures. (A) Gaussian kernel density estimates for affective arousal ratings across an example subset of participants. A distribution reflects the frequency of responses on the affective arousal scale for a particular participant across the experiment. (B) Timeseries of affective arousal ratings across all 50 trials for an example set of 3 participants. Each point reflects a particular affective arousal rating submitted by a participant on one trial. (C) Within-participant posterior distributions of the slope parameter between affective arousal and measures of both physiological arousal and other dimensions of ongoing experience. For experience measures, higher values on the x-axis reflect (i) increased future thinking, (ii) increased deliberate thinking, (iii) increased self-related thinking, and (iv) increased difficulty to disengage from thinking. Points reflect means of participant-level posterior distributions; bands around points reflect 95% credible intervals of the participant-level posterior distribution. Top line of panel text reflects the group-level posterior mean and 95% credible interval for the slope parameter. Bottom line reports the *p* value estimated from a frequentist hierarchical model using the Satterthwaite approximation for degrees of freedom.

We next examined the relationship between subjective and physiological arousal measures (see top row of *Figure 2C*). Group-level 95% credible intervals included zero for all physiological measures: time-on-task (β = −0.14, 95 *CIs* = [−1.34, 1.13], *t*(50.21) = 0.24, *p* = .812), heart rate (β = −1.02, 95 *CIs* = [−3.43, 1.25], *t*(52.62) = 0.78, *p* = .441), alpha power (β = −0.02, 95 *CIs* = [−0.04, 0.01], *t*(48.09) = 1.21, *p* = .232), and HRV (β = −0.003, 95 *CIs* = [−0.01, 4.92 × 10^-1^], *t*(2302) = 1.99, *p* = .047). Note that, while frequentist estimation indicated a significant association between HRV and affective arousal (*p* = .045), the Bayesian 95% credible interval included zero. This discrepancy was likely driven by extremely low between-participant variance in the HRV slope estimate (σ = 0.002; see Figure 2C), which inflated the estimated degrees of freedom and, in turn, lowered the threshold for significance in the *t* distribution. These results converge to support the idea that physiological measures of arousal were largely unrelated to affective arousal ratings at the group-average level.

Having explored the relationship between subjective and physiological arousal, we next sought to more fully characterize what affective arousal measures capture by examining its relationship with other self-reported dimensions of ongoing experience (see bottom row of *Figure 2C*). We report results from hierarchical Bayesian linear regression models for the four experience sampling items that exhibited the highest average participant-level correlations with affective arousal. Group-level 95% credible intervals excluded zero for each of these experience sampling items, indicating significant relationships with affective arousal. Specifically, higher levels of affective arousal were associated with increased future thinking (β = 5.03, 95 *CIs* = [2.89, 7.16], *t*(52.05) = 4.86, *p* < .001), increased deliberate thinking (β = 4.69, 95 *CIs* = [2.54, 6.92], *t*(52.0) = 4.30, *p* < .001), increased self-related thinking (β = 4.45, 95 *CIs* = [2.19, 6.61], *t*(51.49) = 3.97, *p* < .001), and increased difficulty disengaging from thinking (β = 3.51, 95 *CIs* = [1.44, 5.56], *t*(48.62) = 3.34, *p* = .002). While affective arousal was overall related to these four dimensions of ongoing experience, the moderate strength of the standardized regression coefficients across participants suggests that affective arousal carries independent variance from other dimensions of ongoing experience.

### Lower affective arousal is associated with greater HEP amplitude

To assess the relationship between affective arousal and cardiac interoceptive sensitivity, we examined the association between the HEP and affective arousal ratings using a regression-based cluster permutation analysis. This analysis revealed a significant cluster at frontal electrodes (F3, F4, FC1, FC2, Fp1, Fp2, Fz; see *Figure 3B*) from 0.340 to 0.356 s post R peak, where regression coefficients were negative in this cluster (see *Figure 3A*), suggesting that decreased affective arousal ratings are associated with increased HEP (see *Figure 3C*. The direction of this effect was largely consistent across participants (see *Figure 3D*). A surrogate cluster control test revealed that the cluster-wise statistic is significantly greater than what would be expected from a null effect not time-locked to heartbeats (*p* = .02; see *Figure S3*). Since larger HEP amplitude is thought to reflect greater interoception (Judah et al., 2018; Pang et al., 2019; Verdonk et al., 2024), the direction of this effect implies lower affective arousal during states of greater cardiac interoceptive sensitivity. Interestingly, the topology, time course, and direction of this effect closely replicate and extend a finding from past work, where affective arousal was driven by an external event rather than spontaneously fluctuating at rest (Fourcade et al., 2024).

**Figure 3.**
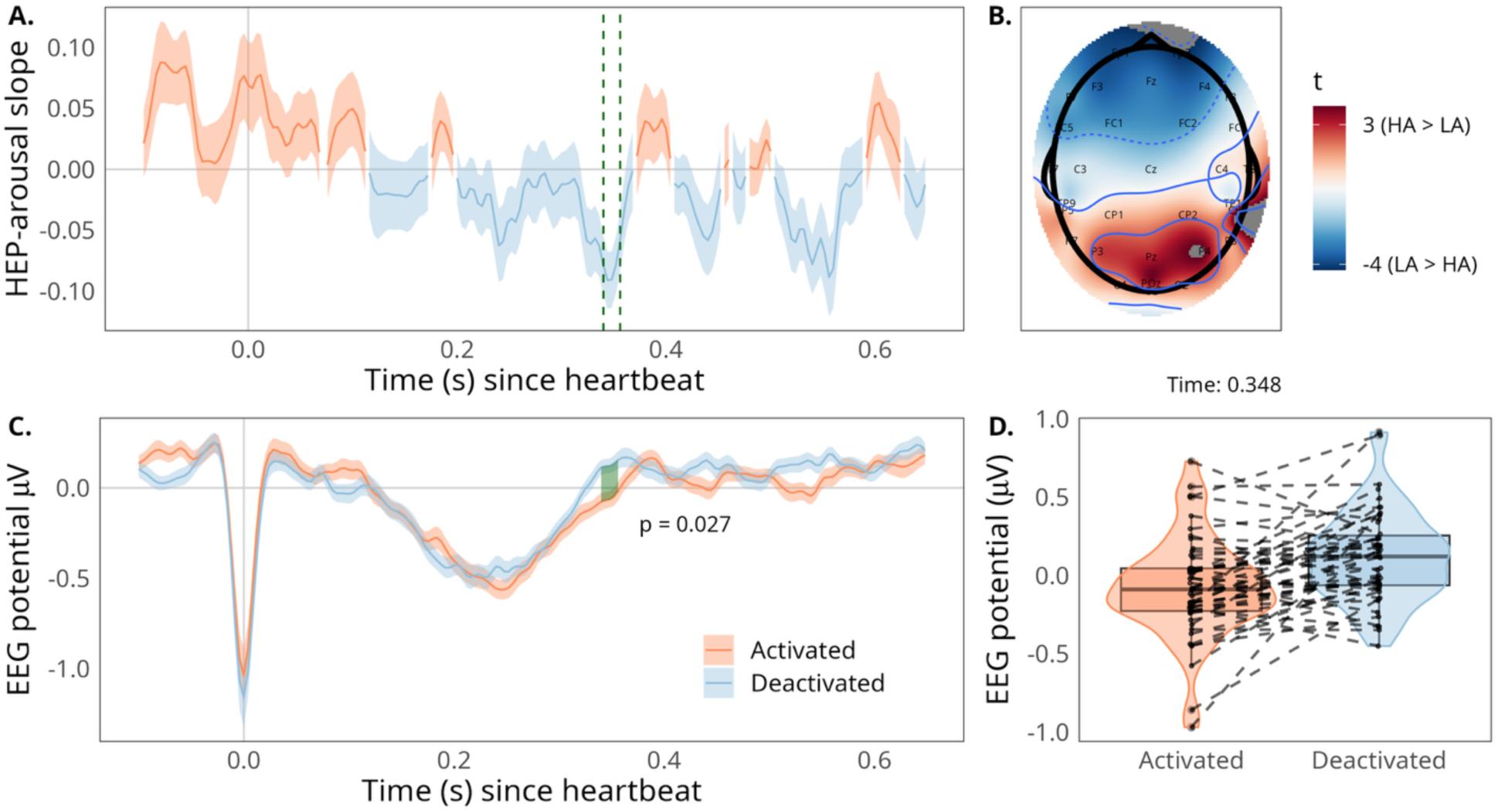
HEP effect across time, participants, and electrodes. (A) Average regression slope across the heartbeat epoch. Timepoint zero reflects the R peak, and colors denote whether EEG voltage was positively (orange) or negatively (blue) associated with affective arousal ratings. Shaded regions reflect standard error of the mean. The green dashed lines reflect the significant time window indicated by the cluster-based permutation analysis. The regression slopes are averaged across significant electrodes. (B) Topographical map of *t*-values for slopes comparing HEP amplitude to affective arousal. (C) HEP grand average across the heartbeat epoch, visualized across within-participant median split in affective arousal ratings. Timepoint zero reflects the R-peak, and colors denote the affective arousal direction. The green shaded region reflects the significant time window indicated by the cluster-based permutation analysis (0.340 – 0.356 s), while the colored shaded regions reflect standard errors of the mean. The EEG waveform is averaged across significant electrodes (i.e., F3, F4, FC1, FC2, Fp1, Fp2, Fz). (D) HEP averaged across a participant-level median split in affective arousal collapsed across the significant time window and channels, shown separately for each participant (dotted lines connect averages across conditions from the same participant).

### The relationship between affective arousal and cardiac interoceptive sensitivity depends on trait anxiety

To investigate whether increased cardiac interoceptive sensitivity during states of lower affective arousal is linked to anxiety, we examined whether the size of the participant-level HEP effect correlated with individual difference measures in both trait (GAD) and state (STAI-S) anxiety. While there was a non-significant relationship between the HEP effect and STAI-S (*r*(29) = −.25, *p* = .182), there was a significant positive correlation between the HEP effect and GAD (*r*(49) = −.34, *p* = .015; see *Figure 4A*). In other words, the extent to which HEP amplitude was more positive for lower states of arousal was increased for those individuals who reported greater trait anxiety (*Figure 4B*).

**Figure 4.**
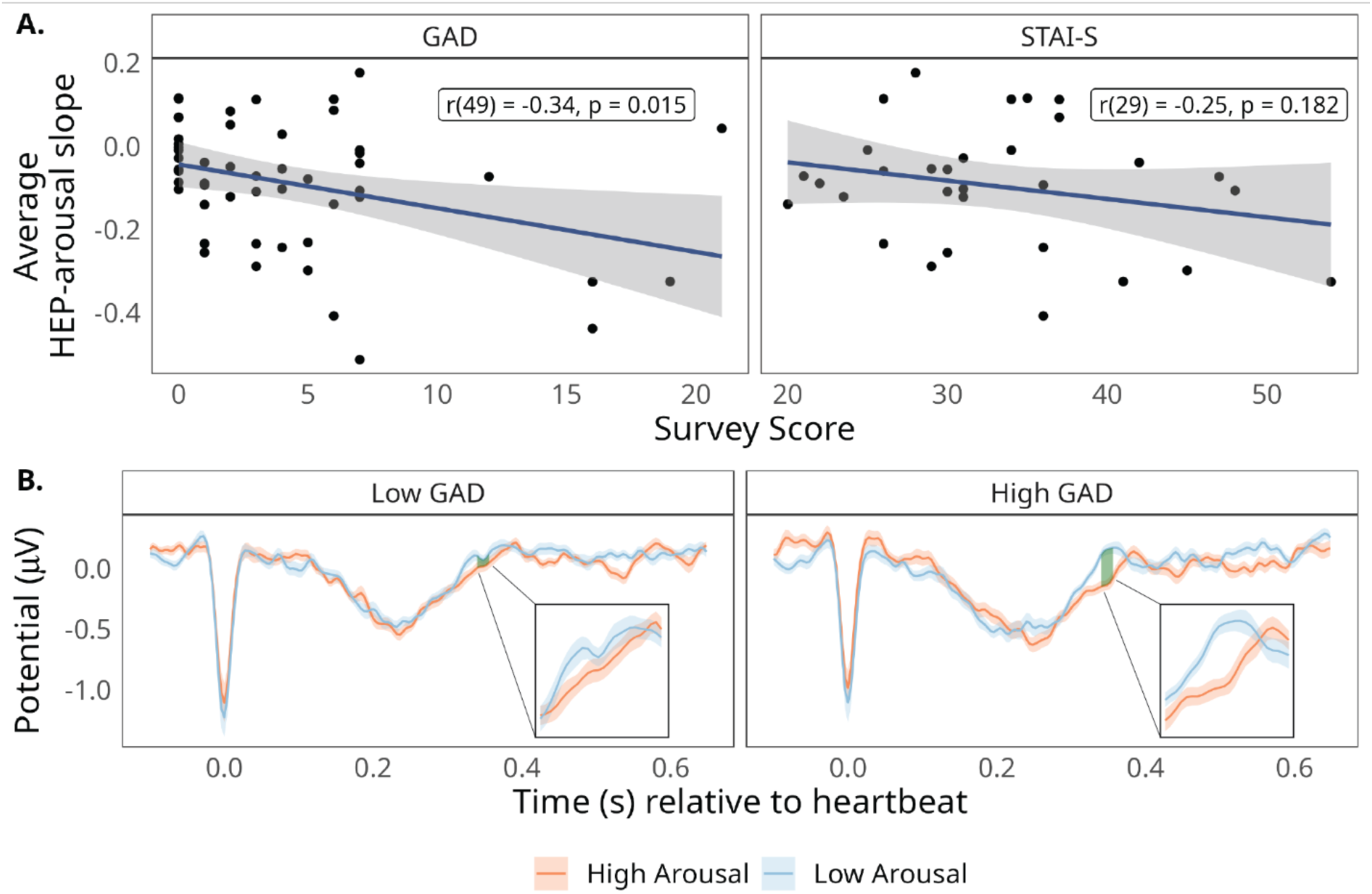
Association between HEP arousal effect (averaged HEP-arousal slope) and individual differences in anxiety. (A) Correlations of trait (GAD) and state (STAI-S) anxiety with individual differences in the HEP effect magnitude (lower on the y-axis reflects larger HEP amplitude for low vs. high affective arousal). (B) HEP grand average across the heartbeat epoch, broken down by within-participant median split on affective arousal ratings and a median split on trait anxiety (GAD). Insets zoom in to the significant time window, indicated by the shaded green region. Shaded colored regions reflect one standard error from the mean.

## Discussion

Our experiment found that cardiac interoceptive sensitivity increased during lower affective arousal during spontaneous thought, with this effect amplified in individuals with high trait anxiety. Affective arousal was largely distinct from physiological measures like time-on-task, parieto-occipital alpha power, and heart rate, though the relationship with heart rate variability was more equivocal. Affective arousal also differed from other aspects of spontaneous mental experience and was positively associated with future thinking, deliberate thinking, self-related thinking, and difficulty disengaging from thoughts. Cardiac interoceptive sensitivity, indexed by the HEP, showed greater amplitude prior to lower affective arousal ratings, localized at frontal electrodes (0.340 – 0.356 ms post-heartbeat). This effect was stronger in those with high trait anxiety. Our findings refine the concept of arousal and have clinical relevance for anxiety and interoceptive dysregulation (Khalsa & Lapidus, 2016).

Arousal is widely studied but inconsistently defined (Lindquist et al., 2016; Satpute et al., 2019). Measures such as self-report, skin conductance, heart rate, respiratory frequency, and EEG alpha power often show weak or inconsistent correlations among each other (Agren, 2023; Leonidou & Panayiotou, 2022; Wenzler et al., 2017). This inconsistency has led some to conceptualize arousal as multidimensional (Reid et al., 2024), while others argue that distinct types of arousal may share underlying cortical pathways, namely the presupplementary motor areas and dorsal anterior insula (Sabat et al., 2025). These competing perspectives underscore the need to differentiate aspects of arousal. In our data, spontaneous fluctuations in self-reported affective arousal showed little association with physiological measures: 95% credible intervals for all physiological predictors included zero. Although HRV reached nominal significance in a frequentist model, the effect size was small and exhibited minimal between-subject variability. These findings add to growing evidence that subjective and physiological arousal may reflect distinct components of a broader arousal construct.

Our findings lend support to the idea that arousal is a multidimensional construct with independent subcomponents (Reid et al., 2024). Although arousal has been widely studied, few investigations have examined spontaneous fluctuations in affective arousal during wakeful rest (but see Hoemann et al., 2020). The association between affective and physiological arousal may depend on whether attention is focused on external stimuli (Leonidou & Panayiotou, 2022), raising the possibility that self-reported arousal reflects deliberate rather than purely physiological processes. Future research should examine how spontaneous fluctuations in affective arousal relate to attentional focus and how this interaction influences the link (or lack thereof) between subjective and physiological arousal.

Affective arousal was moderately associated with other dimensions of ongoing experience, such as future thinking, deliberate thinking, self-related thinking, and difficulty to disengage from thoughts. Deliberate thinking, which involves effortful problem solving, has been linked to increased physiological and affective arousal, particularly when deliberation is framed as task engagement (Kahneman, 2011; Matthews et al., 2010). Our results extend these ideas by showing a positive link between affective arousal and deliberate thinking even when deliberate thinking is not tied to an active task. Difficulty disengaging from thoughts—often linked to repetitive negative thinking and worrying, key features of generalized anxiety disorder (Ehring & Watkins, 2008) and state anxiety (Vălenaş et al., 2017)—was also associated with affective arousal. This finding aligns with prior research showing that self-reported trait anxiety positively correlates with physiological arousal (Hoehn-Saric & McLeod, 2000; Noteboom et al., 2001; Roos et al., 2021). Finally, affective arousal was positively associated with both self- and future-thinking. This finding contrasts somewhat with recent work showing a negative association, at the inter-individual level, between conceptual thinking (as opposed to bodily focus) and physiological arousal (Banellis et al., 2024). Self- and future-thinking tend to co-occur and are associated with improved future mood (Ruby et al., 2013). One possibility is that affective arousal facilitates this relationship, where engaging in self- and future-thinking enhances arousal, which in turn supports mood regulation. However, how affective arousal and self- and future-thinking impact future mood likely depends on individual differences in anxiety. Anxiety has been associated with more vivid and detailed autobiographical future thinking (Du et al., 2022), which may amplify or distort the impact of future thinking on mood.

We investigated how interoceptive processing varies as a function of spontaneous fluctuations in affective arousal by examining the HEP. We found that in the 340–356 ms window post-heartbeat, HEP amplitude was increased at frontal electrodes in the 10 s preceding lower affective arousal ratings. This finding suggests that interoceptive processing is enhanced during states of lower affective arousal. The timing and topography of this effect align with previous studies linking the HEP to arousal (Fourcade et al., 2024; Fukushima et al., 2011; Gentsch et al., 2019; Ito et al., 2019; Luft & Bhattacharya, 2015; MacKinnon et al., 2013; Marshall et al., 2017, 2018, 2020; Park et al., 2016; Schulz et al., 2013, 2015; Sel et al., 2017; Shao et al., 2011), though definitions of arousal vary widely across studies. One study particularly relevant to our work recorded electrophysiological responses while participants experienced virtual reality (VR) roller coaster rides and retrospectively rated fluctuations in affective arousal (Fourcade et al., 2024). Their results closely parallel ours, showing increased HEP for lower affective arousal at fronto-central electrodes in a similar time window. This convergence across methodologies strengthens the link between affective arousal and interoceptive processing indexed by the HEP. Our findings extend this prior work that showed differences in HEP driven by external events (i.e., roller coaster videos; Fourcade et al., 2024) by revealing that such differences in HEP can also be driven by spontaneously fluctuating internal states during wakeful rest. Fourcade et al. (2024) performed source localization analyses on their EEG results to suggest that their effect, which closely parallels ours, originates from somatosensory areas. Indeed, past work using intracranial EEG has implicated a variety of brain regions in HEP activity during rest, including somatosensory cortex, insula, opercular cortex, inferior frontal gyrus, and amygdala (Babo-Rebelo, Wolpert, et al., 2016; Canales-Johnson et al., 2015; Kern et al., 2013; Park et al., 2018; Park & Blanke, 2019).

One possible explanation for increased HEP during lower affective arousal is that reduced arousal may, in some contexts, enhance interoception. This inverse relationship is evident in practices like meditation, which both lower arousal and improve interoceptive awareness (Hanh, 2015; Kushner & Marnocha, 2008; Nathoo, 2021; Shapiro & Carlson, 2009). In Theravada Buddhism, bodily calm is cultivated to heighten sensitivity to internal sensations and support insight into impermanence (Analayo, 2018). Empirical work links meditation to enhanced interoceptive awareness, potentially through altered neural structure and function, particularly in the insula (Farb et al., 2013; Hölzel et al., 2008; Luders et al., 2012). An alternative explanation is that lower HEP during high arousal reflects a compensatory downregulation of interoception— especially in individuals prone to anxiety, who may suppress bodily awareness during high-arousal states (Russell, 1980, 2003). Supporting this view, the present study found larger HEP differences among individuals higher in trait anxiety (see orange lines, Figure 4B). Although clinical anxiety has been linked to disrupted interoception, findings remain mixed (Abrams et al., 2018; Ardizzi & Ferri, 2018; Gaebler et al., 2013; Khalsa & Lapidus, 2016; Stewart et al., 2001). These results underscore the importance of considering how momentary affective arousal shapes the interplay between anxiety and interoceptive processing.

This study had several limitations that can be addressed in future work. The state anxiety measure (i.e., STAI-S) was a late addition to the experiment, and as such we have data from only 30 of the 51 total participants. Our sample also contained few individuals scoring high on trait anxiety (i.e., GAD; see *Figure 4a*). The present experiment was not specifically designed to examine these correlations and may have been underpowered to detect them; thus, associations with anxiety measures should be interpreted as preliminary. Future studies should replicate our findings related to trait anxiety in a larger and more clinically diverse sample. Such a replication is particularly important given the inconsistent links between HEP and anxiety in the literature (Qi et al., 2025). A further limitation concerns the degree to which findings in our laboratory setting translate to real-world contexts. During daily life, many individuals are rarely truly at rest (i.e., devoid of prominent external stimuli or task demands). What may prove crucial in the study of mental health is understanding how spontaneously fluctuating internal states of affective arousal interact with salient external conditions or environmental cues to influence the regulation of anxiety.

In conclusion, our findings offer advanced knowledge of the neural basis of affective arousal and have broad implications for understanding various mental health conditions marked by emotional and interoceptive dysregulation. The interactions between HEP, affective arousal, and anxiety that we identified suggest a potential naturally-operating mechanism in which interoception is spontaneously downregulated to facilitate emotional regulation. Further research is needed to causally test for such a mechanism, how it may be modulated, and whether evidence of this mechanism is present in other clinical populations, such as individuals with generalized anxiety disorder.

## Code Accessibility

We will make the custom code used to analyze the electrophysiological data available upon publication.

## Acknowledgments

This work was supported by the National Institute of Mental Health of the National Institutes of Health under award numbers R21MH129630 and R21MH127384 (to AK).^1^

## Footnotes

1 The authors declare no competing financial interests.

