## Supplementary material for "Neural sensitivity to the heartbeat is modulated by fluctuations in affective arousal during spontaneous thought": Figure S1

*Contrasting mean ECG amplitude across participant-specific median split in affective arousal.*

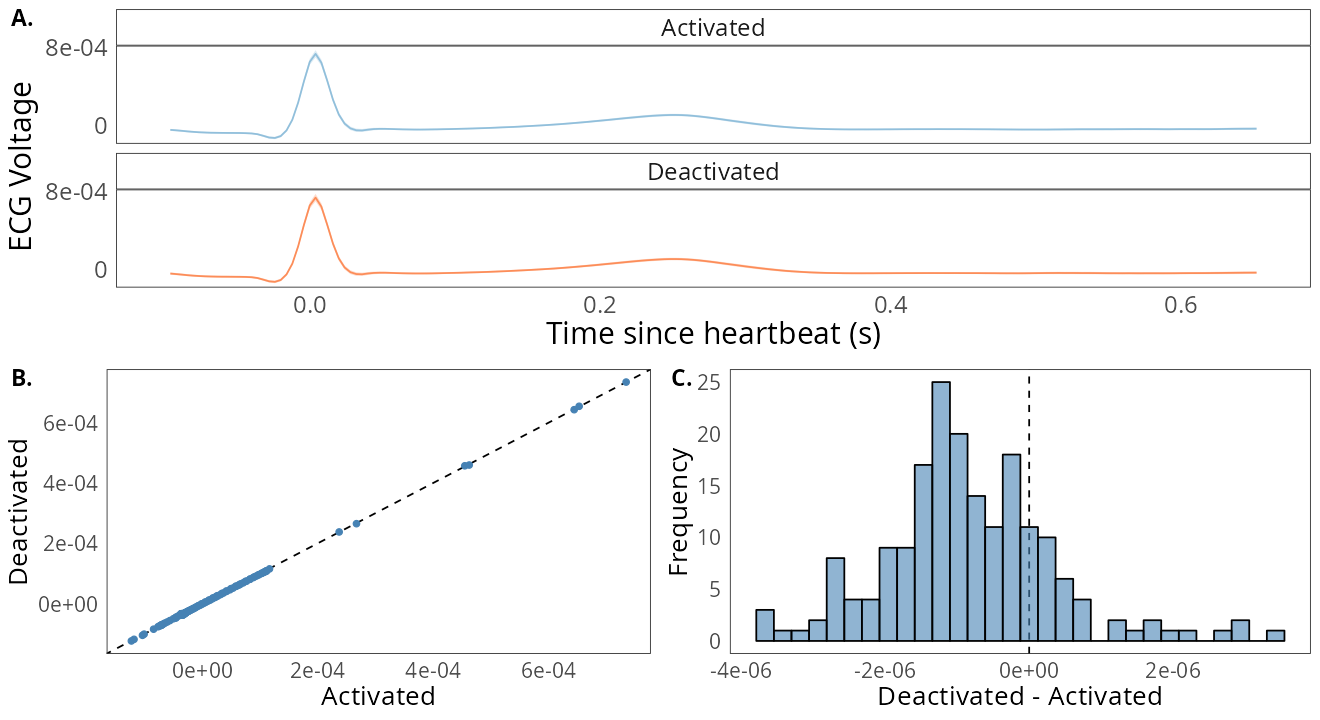
*Note.* The ECG signal was segmented using the same heartbeat epoch window as in the EEG analyses (i.e., −1 to 650 ms around heartbeats), extracted from the 10 s pre-probe window, and averaged within each participant after splitting trials by their median affective arousal rating, then grand averaged across participants. Only minimal differences in ECG amplitude were observed between affective arousal conditions (Panels A & B). A histogram of participant-level ECG amplitude differences between low (“Deactivated”) and high (“Activated”) affective arousal reveals a roughly normal distribution centered near zero, indicating no systematic ECG amplitude differences across conditions (Panel C).
