## Supplementary material for "Neural sensitivity to the heartbeat is modulated by fluctuations in affective arousal during spontaneous thought": Figure S2

*Subjective Arousal Distributions by Participant*

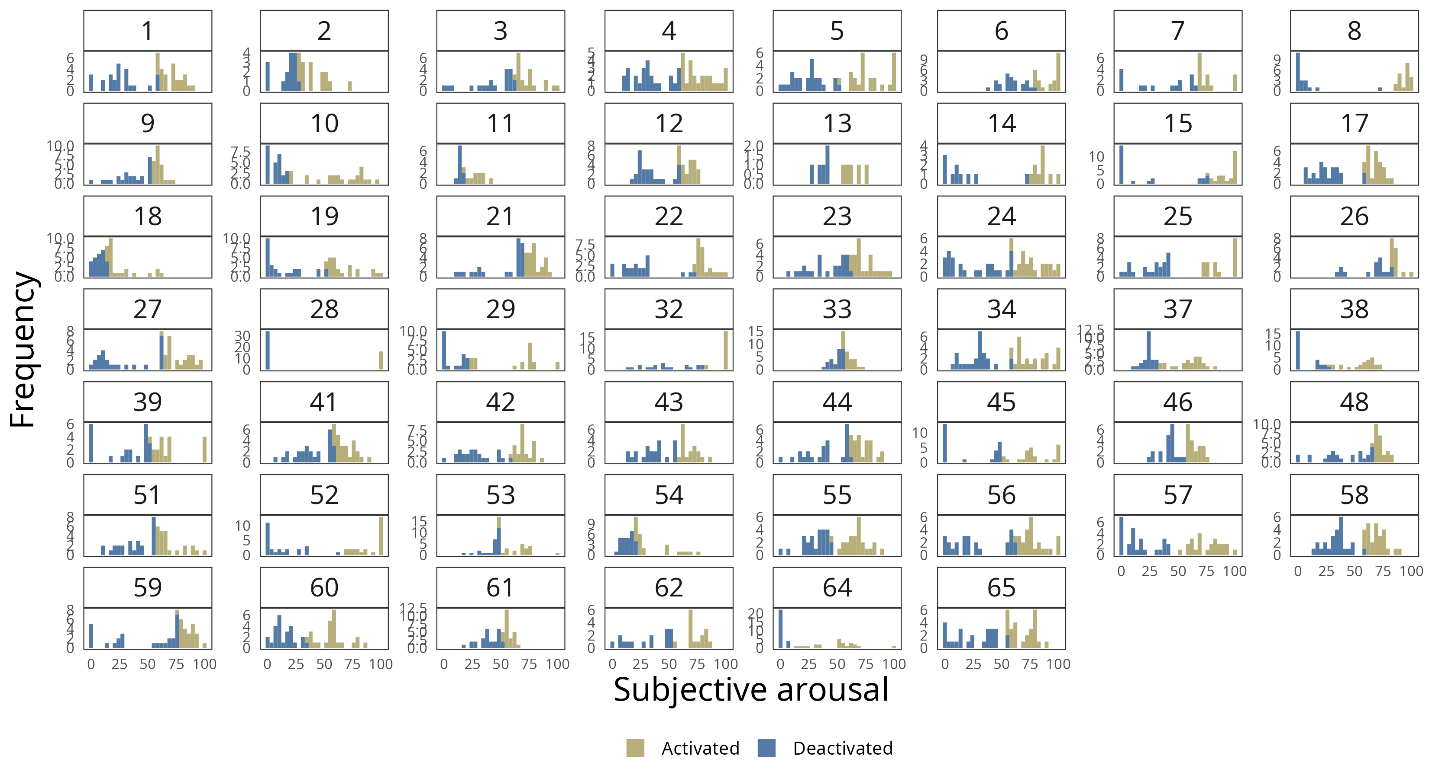

*Note.* Histograms reflect the observed distribution of affective arousal ratings (0 – 100) colored by a median split for each participant over the entire experiment (“Activated” maps to high affective arousal; “Deactivated” maps to low affective arousal).
