## Supplementary material for "Neural sensitivity to the heartbeat is modulated by fluctuations in affective arousal during spontaneous thought": Figure S3

*Results of surrogate cluster control test*

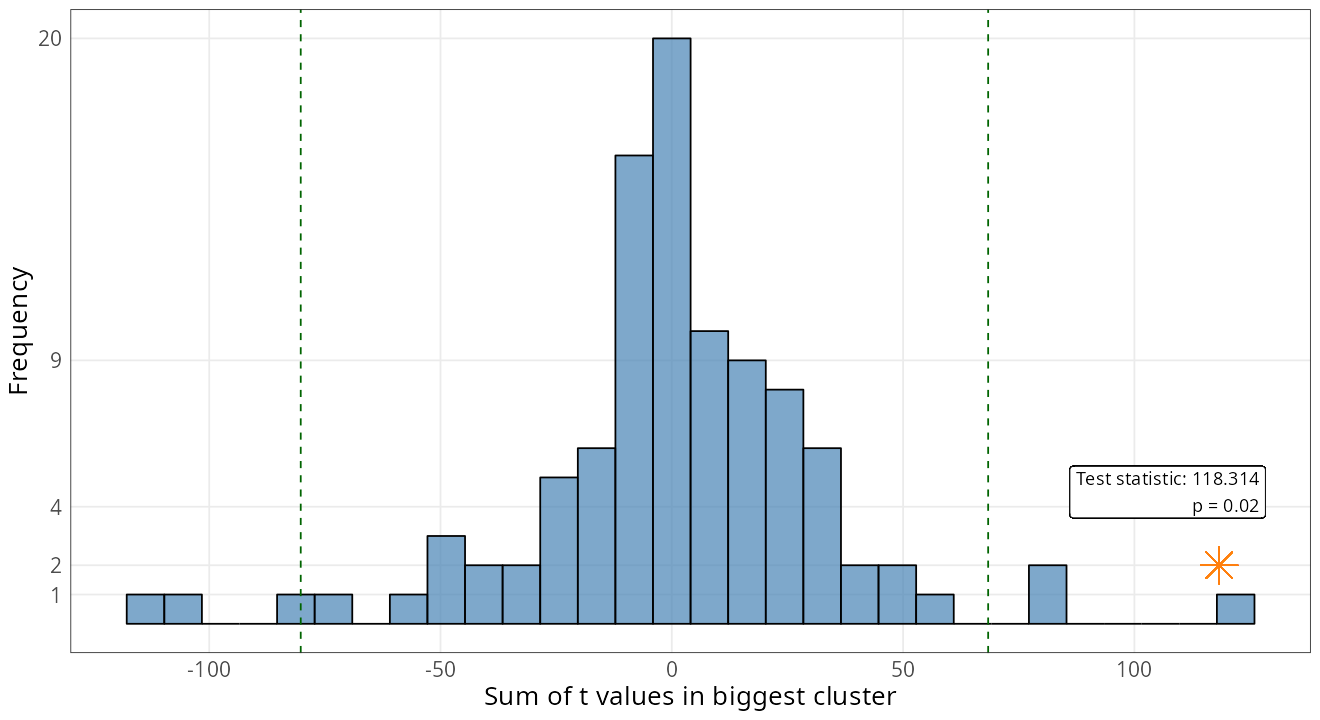

*Note.* The summed *t* value derived from summarizing epochs time-locked to heartbeats (118.314) was significantly greater (*p* = .02) than a null distribution of summed *t* values obtained from summarizing epochs that were randomly shuffled in time (i.e., decoupled from heartbeats).
