## Supplementary material for "Neural sensitivity to the heartbeat is modulated by fluctuations in affective arousal during spontaneous thought": Table S1

*Mental Health Surveys*

| Category | Scale name | Reference |
| --- | --- | --- |
| Crosscutting mental health assessment | DSM-5 Self-Rated Level 1 Cross-Cutting Symptom Measure-Adult (DSM-XC) | American Psychiatric Association, 2013 |
| Measure of impairment | WHO Disability Assessment Schedule-12 (WHODAS-12) | Üstün, 2010 |
| Anxiety and depression measures | Patient Health Questionnaire-9 (PHQ-9) | Kroenke et al., 2001 |
|  | Ruminative Response Scale (RRS) | Treynor et al., 2003 |
|  | General Anxiety Disorder-7 (GAD-7) | Spitzer et al., 2006 |
|  | State Trait Anxiety Inventory (state scales; STAI-S) | Speilberger et al., 1983 |
| Mind wandering | Mind-Wandering Deliberate-Spontaneous (MWD-S) | Carriere et al., 2013 |

*Note.* A collection of mental health surveys, all of which participants were asked to complete prior to arriving to the experiment, except for the state scale of the State Trait Anxiety Inventory (STAI-S), which was completed at the beginning of the experimental session.
