## Supplementary material for "Neural sensitivity to the heartbeat is modulated by fluctuations in affective arousal during spontaneous thought": Table S2

*Complete Set of Experience Sampling Items presented in each Thought Probe*

| Variable | Question | Description | Low anchor (0) | High anchor (100) | |
| --- | --- | --- | --- | --- | --- |
| Attentional focus | Were you more focused on your thoughts (mental) or sensing the world or your body (physical)? (or if your mind was blank press the 0 key) | Determine whether your attention was on your thoughts (mental) or something tangible on your body or the environment (physical) or if you are unable to recall thinking about anything (mind blank) | Completely physical | | Completely mental |
| Past | Were your thoughts oriented towards the past? | Determine where your thoughts were located in time | Not past oriented | | Completely past oriented |
| Future | Were your thoughts oriented towards the future? | Determine where your thoughts were located in time | Not future oriented | | Completely future oriented |
| Self | Were your thoughts about yourself? | Think about whether your thoughts were focused on yourself, or matters related to you | Nothing about you | | Completely about you |
| Others | Were your thoughts about others? | Think about whether your thoughts were focused on yourself, or matters related to you | Nothing about others | | Completely about others |
| Arousal | How activated or energized were you feeling? | Activation (or arousal) includes feelings of energy linked to emotion. | Completely deactivated | | Completely activated |
| Affect | How positive or negative were you feeling? | How positive or negative you felt while having these thoughts | Completely negative | | Completely positive |
| Disengage | How easy was it to disengage from your thoughts? | Think about how easy it was to pull yourself away from your thoughts at the onset of the probe | Extremely easy | | Extremely difficult |
| Movement | Were your thoughts freely moving? | Think about the last few seconds before the questions popped up and whether you were focused on one thought for an extended period of time (unmoving) or if your thoughts were drifting from one thing to another without focusing on any single thought for long (freely moving). | Unmoving | | Moving freely |
| Intentional | How intentional were your thoughts? | Think about whether you directed your attention to a certain task or thoughts with purpose or if your thoughts popped into your mind without intending to do so. | Completely unintentional | | Completely intentional |
| Visual | Were your thoughts visual? | Think about whether your thoughts took the form of images | Completely visual | | Completely non-visual |
| Verbal | Were your thoughts verbal | Were your thoughts manifesting as a voice in your head or were your thoughts oriented as words or using language. | Completely verbal | | Completely non-verbal |
| Confidence | How confident are you about your ratings for this trial? | How accurately you were able to remember and categorize your thoughts by answering the questions, and how strongly you believe they reflect your last thought. | Completely confident | | Completely unconfident |

*Note*. For each experience sampling item, the text under the “Question” column was displayed on the top of the screen, and on the bottom of the screen was a response slider with the text in the “Low anchor (0)” column displayed on the left and the text in the “High anchor (100)” column displayed on the right. Experimenters recited the text under the “Definition” column to participants during the instructional phase of the experiment.
